# Are working memory training effects paradigm-specific?

**DOI:** 10.1101/450023

**Authors:** Joni Holmes, Francesca Woolgar, Adam Hampshire, Susan E. Gathercole

**Affiliations:** MRC Cognition & Brain Sciences Unit, University of Cambridge, UK; Imperial College London, UK; University of Cambridge, UK

**Keywords:** working memory, cognitive training, transfer, intervention, memory

## Abstract

A randomized controlled trial compared complex span and n-back training regimes to investigate the generality of training benefits across materials and paradigms. The memory items and training intensities were equated across programs, providing the first like-with-like comparison of transfer in these two widely-used training paradigms. The stimuli in transfer tests of verbal and visuo-spatial n-back and complex span differed from the trained tasks, but were matched across the untrained paradigms. Pre-to-post changes were observed for untrained n-back tasks following n-back training. Following complex span training there was equivocal evidence for improvements on a verbal complex span task, but no evidence for changes on an untrained visuo-spatial complex span activity. Relative to a no intervention group, the evidence supported no change on an untrained verbal complex span task following either n-back or complex span training. Equivocal evidence was found for improvements on visuo-spatial complex span and verbal and visuo-spatial n-back tasks following both training regimes. Evidence for selective transfer (comparing the two active training groups) was only found for an untrained visuo-spatial n-back task following n-back training. There was no evidence for cross-paradigm transfer. Thus transfer is constrained by working memory paradigm and the nature of individual processes executed within complex span tasks. However, within-paradigm transfer can occur when the change is limited to stimulus category, at least for n-back.

## Introduction

Research investigating transfer within and across working memory paradigms following training has largely concluded that transfer is confined to untrained working memory tests that are highly similar to the trained activities (e.g. Blacker, Negooita, Ewen & Courtney, 2017; Harrison, Shipstead, Hicks, Hambrick, Redick & Engle, 2013; Minear, Brasher, Guerro, Brasher, Moore & Sukeena, 2016; Sprenger et al., 2013; Soveri, Antfolk, Karlsson, Salo & Laine, 2018; von Bastian, Langer, Jancke & Oberauer, 2013). This has led to speculation that training induces changes in the processes and strategies tied to particular training experiences rather than fundamental improvements in the more general capacity of the working memory system (see Dahlin, Neely, Larsson, Backmann & Nyberg, 2008; Dunning & Holmes, 2014; Sprenger et al., 2013; von Bastian & Oberauer, 2013, 2014). Progress in understanding the boundary conditions for transfer and the nature of the cognitive training changes induced by training has been hampered by the absence of robust theories of transfer and, consequently, a lack of hypothesis-driven research. Experimental studies manipulating the overlap in key task features between training and transfer measures are needed to identify the precise conditions under which transfer does and does not occur (e.g. Minear et al., 2016; von Bastian et al., 2013). The current study addressed this by examining the degree to which near transfer across working memory tasks is limited by two properties of both the trained and untrained tasks: the stimulus category and the working memory paradigm.

Several process- and task-specific theories of transfer have been advanced. One proposal is that the efficiency of specific kinds of working memory activity may be enhanced through training. Dahlin et al. (2008), for example, suggested that training on n-back tasks that require the continuous updating of the contents of working memory enhances a process of updating the contents of working memory that can be applied to other updating paradigms such as running span. Gathercole, Dunning, Holmes & Norris (2018) adopted a broader perspective and speculated that training leads to the development of paradigm-specific skill-like cognitive routines when the working memory task is an unfamiliar one that is not readily supported by the basic short-term memory system. This theory predicts that transfer will be limited by paradigm: a cognitive routine that is learned through training will only benefit untrained tasks with the same higher-level cognitive structure. Alternatively, training may lead to the development of explicit mnemonic strategies such as learning to group individual memory items into larger chunks, or recoding items into alternative and more distinctive forms (Dunning & Holmes, 2014; Minear et al., 2016; von Bastian & Oberauer, 2013). A further possibility is that transfer may not have a single origin at all but is instead the product of multiple training-induced changes in processes specific to individual task features. These could include the nature of stimulus items, the response modality, and the timing parameters of the task, as well as the broader paradigms in which they are embedded (Minear et al.,2016; Sprenger et al., 2013; von Bastian & Oberauer, 2013).

The current study examined the limits to transfer in two commonly-used training regimes, complex span and n-back. Transfer to untrained n-back tasks following n-back training is common (e.g. Soveri et al., 2017). This holds true when the stimulus items are either from both the same (e.g. letters to digits) or different domain (e.g spatial locations to digits) to the trained items (Anguera et al., 2012; Blacker et al., 2017; Bürki, Ludwig, Chicherio & de Ribaupierre, 2014; Jaeggi, Buschkuehl, Perrig & Meier, 2010; Li. Schmiedek, Huxhold, Röcke, Smith, & Lindenberger, 2008; Minear et al., 2016; Küper & Karbach, 2016; Waris, Soveri & Laine, 2015). All complex span training studies to date have changed both the storage items and the nature of the interpolated distraction (e.g. symmetry judgements vs spatial orientation decisions, or sentence judgements vs mathematical operations) between trained and untrained tasks. Significant transfer has been reported across the majority of these studies (Harrison et al., 2013; Henry, Messer & Nash, 2014; von Bastian et al., 2013; Richmond, Wolk, Chein & Morrison, 2010). However, transfer across stimulus categories is not always consistent. Minear et al. (2016) reported mixed patterns of transfer within and across domains in different conditions. Following training on a verbal complex span task transfer was found for operation and rotation span, but not for symmetry or alignment span. Blacker et al. (2017) reported no transfer between two verbal variants of complex span.

The evidence to date largely indicates little transfer across n-back and complex span paradigms. Although a few studies report positive transfer following n-back training to a variety of complex span tasks (Anguera et al., 2012; Minear et al., 2016; Schwarb, Nail & Schumacher, 2015; Sprenger et al., 2013), the majority have failed to observe transfer (Blacker et al., 2017; Burki et al., 2014; Chooi & Thompson, 2012; Li et al., 2008; Thompson et al., 2013; Jaeggi, Buschkeuhl, Jonides & Perrig, 2008; Jaeggi et al., 2010; Lilienthal, Tamez, Shelton, Myerson & Hale, 2013; Minear et al., 2016; Redick et al., 2013; Schwarb et al., 2015; Sprenger et al., 2013). Only three studies have tested transfer from complex span training to n-back. Minear et al. (2016) reported transfer to an object n-back task but not to a letter n-back task following training on a verbal complex span task. von Bastian et al. (2013) failed to find transfer to a letter n-back task following training on a verbal complex span task. Blacker et al. (2017) found that training on a symmetry span task did not improve performance on an object n-back task.

The primary purpose of the study was to test whether transfer occurs with a paradigm when the stimuli between training and transfer (for both paradigms) and the interpolated distractor activity (for complex span) differ. It is the first to compare the two training approaches while matching potentially key features for transfer across paradigms: the stimuli in the transfer tests within and across paradigm, and the duration of training of the two training regimes. Holding these features constant provides a direct test of whether paradigm is the critical factor constraining transfer.

It was predicted that transfer would be limited to new tasks employing the same working memory paradigm. The weight of evidence to date suggests that it is, and this is consistent with accounts of paradigm-specific accounts of transfer (Gathercole et al., 2018). No transfer is therefore predicted between the two paradigms. Minear et al. (2016) is one of two studies to compare directly these two different forms of training. They used spatial n-back and verbal complex span training tasks, and found no evidence for cross-paradigm transfer. However, as both the stimulus category and domain of the memory items differed between the training regimes it is impossible to determine from this study whether transfer was restricted by working memory paradigm or differences in the training stimuli. A second study comparing dual n-back training with locations and letters to a symmetry span training regime, which also failed to find cross-paradigm transfer, is limited by the same confounds; one training regime targeted only visuo-spatial abilities while the other included both verbal and visuo-spatial materials (Blacker et al., 2017). Matching the stimuli across the training paradigms and in the transfer tests enabled us to test the extent to which training paradigm *per se* is a boundary condition to transfer.

Transfer across n-back tasks that differ only in the stimulus category within a common domain was expected on the basis of previous findings (e.g. Jaeggi et al., 2010& Minear et al., 2016; Küper & Karbach, 2016; Soveri et al., 2017). A final question addressed in this study was whether transfer occurs across complex span tasks with different storage items and distractor activities. We tested this by employing trained and untrained complex span tasks with distinct distractor activities (lexical decision and rhyme judgement; symmetry judgement and mental rotation) and memory items (digits and letters; spatial locations and visual objects). Previous evidence has pointed to transfer across tasks sharing only the paradigm and not either the stimuli or distractor activity (Harrison et al., 2013; Henry et al., 2013; Minear et al., 2016; Richmond et al., 2014; von Bastian et al., 2013). The cognitive routine theory (Gathercole et al., 2018) makes no strong predictions about the level of specificity of a complex span routine. If it was a high-level specification reflecting the alternating stimulus presentation and distractor processing episodes, it might be expected to be readily applied to other complex span tasks with different stimuli and distractor activities. Alternatively, the component processes involved in the routine might be specified at a greater level of detail including subroutines for specific distractor activities, in which case no transfer across different complex span tasks would be expected. Either way, the study will provide key information regarding the boundary conditions to transfer that will inform new theory.

**Table 1.**
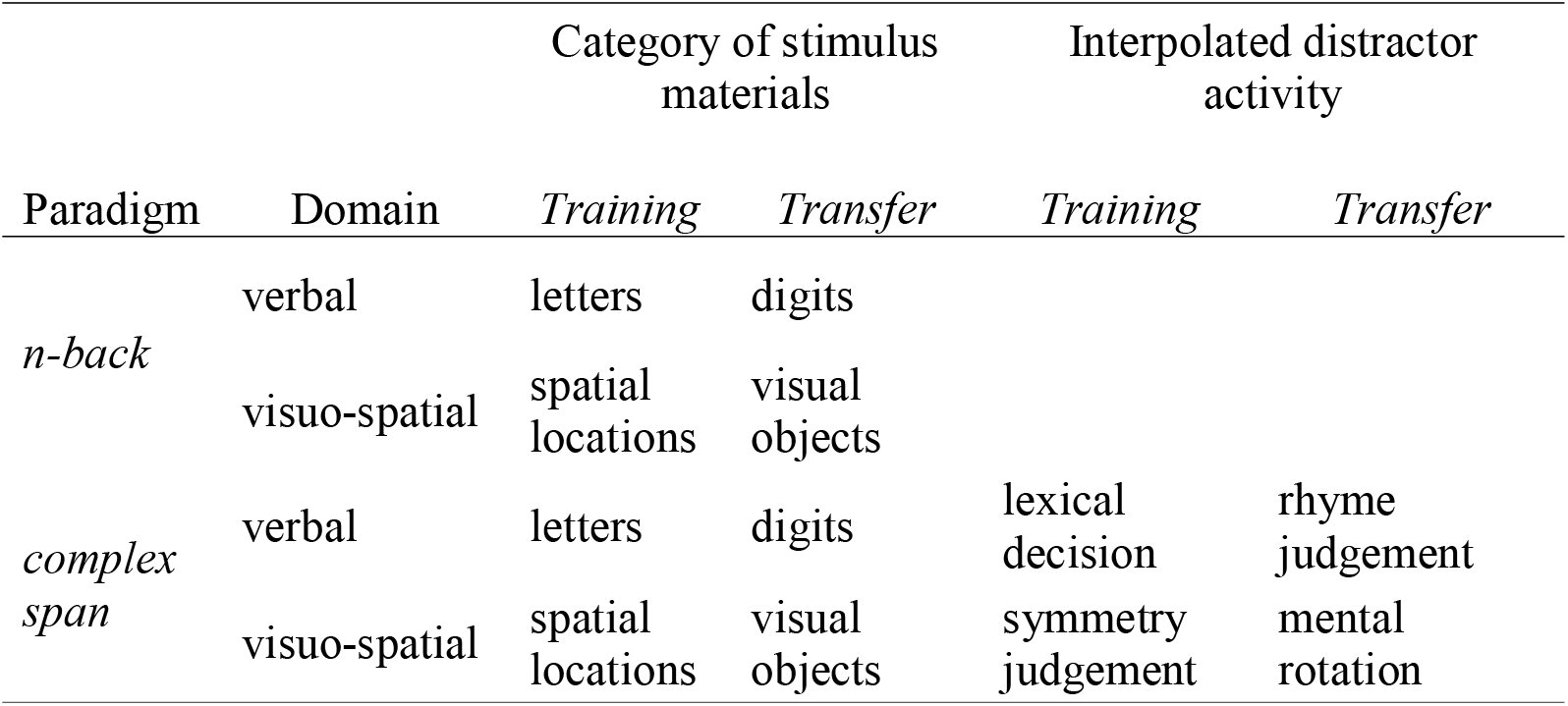
Trained and untrained tasks

The trained and untrained tasks are summarised in Table 1, with images of the tasks shown in Figure 1. Bayesian inference was used to evaluate the strength of the evidence of training and transfer effects. Bayes factors (BF) quantify the evidence for both null hypotheses (the absence of training and transfer) and alternative hypotheses (the presence of training and transfer). BFs are increasingly popular in cognitive training research as a means of quantifying positive evidence for the null hypothesis of no transfer (e.g. DiSimoni & von Bastian, 2018; Sprenger et al., 2013).

**Figure 1.**
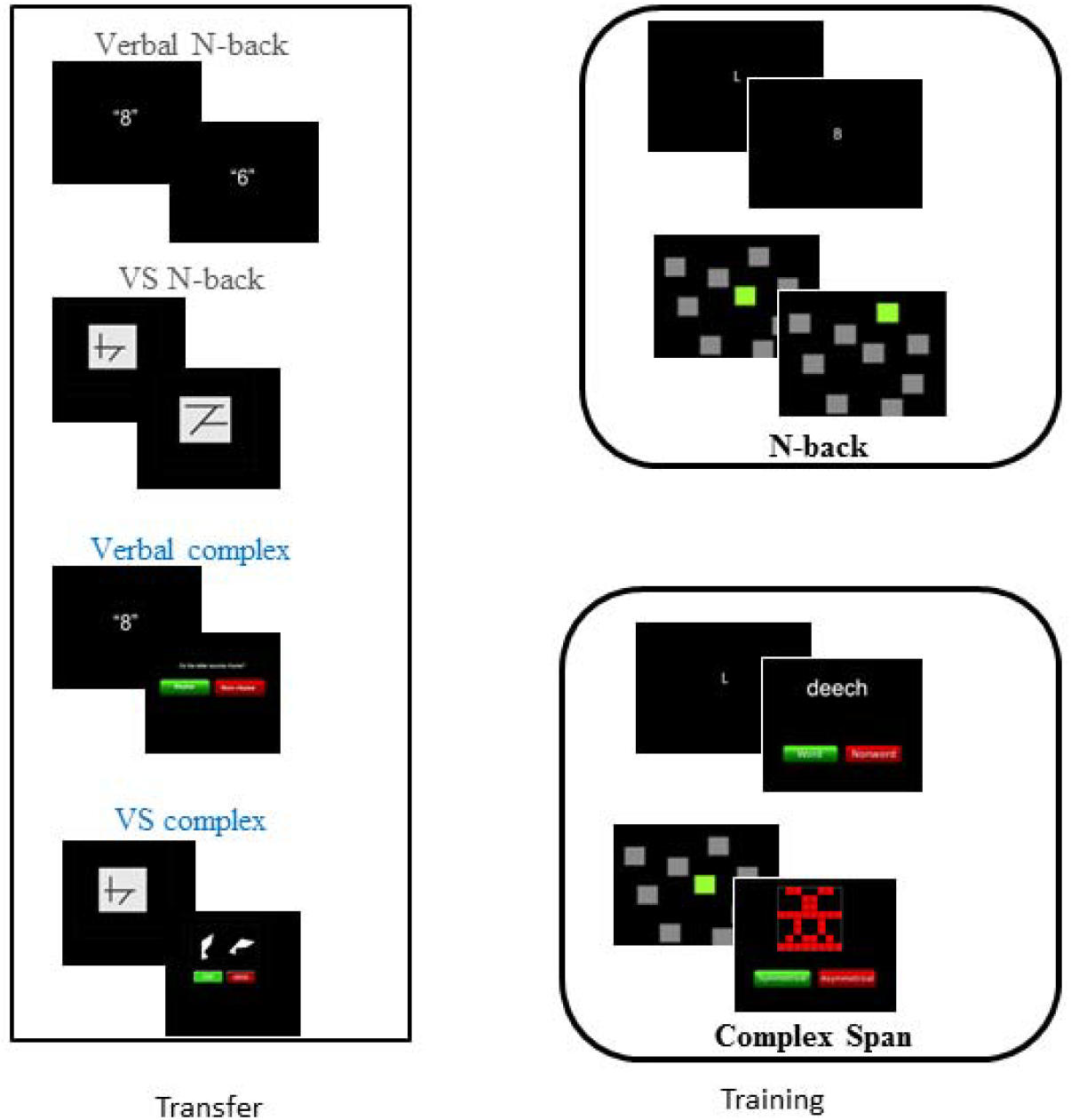
Training and transfer tasks

## Materials and Method

### Participants

Target recruitment was 16 participants per group (total N=48), yielding power of .86 to detect a large effect size, *f^2^*=.35 or Cohen’s d=.8, with linear regression. Data were excluded for eight participants who failed to complete all training sessions and four participants who did not attend for the Time 2 assessment. The training program failed for one participant. Further participants were recruited to replace those with incomplete data.

The final sample included 48 native-English speaking adults aged between 18 and 35 years (12 males, mean age 28 years and 9 months). Participants were recruited through the MRC Cognition & Brain Sciences Unit research participation system and provided written informed consent prior to participation. Ethical approval was obtained from the University of Cambridge’s Psychology Research Ethics Committee (PRE.2012.86).

### Materials

#### Transfer tasks

##### Complex span

Participants completed two complex span tasks, one verbal and one visuo-spatial. For both tasks participants were presented with a series of storage items interpolated with a same-domain processing task performed for a 6000ms period intervening between the presentation of successive memory items. Participants were required to recall the storage items in serial order at the end of the trial. Two practice trials were presented at a list length of one item. Test trials were presented in blocks of 3. The first block started at a list length of one (a single memory item followed by a processing episode prior to recall) and increased by one item (additional storage item and an additional processing episode) if two or more trials were correct in any block. Trials were scored as correct if all storage items were recalled in the correct serial order and >66% of the processing items were correct. The tasks discontinued if two of the three trials in a block were incorrect. A trial was incorrect if the storage items were recalled incorrectly, accuracy for the processing tasks was <66%, or if there were no responses for the processing tasks. The maximum span reached was scored. This was counted as the span level at which the task discontinued.

The auditory stimulus items in the verbal complex span task were the digits 1 through to 9. Digital recordings of each item spoken in a female voice were p-centered - a process of aligning the waveforms of the recordings so the digits sound regular (Morton, Marcus & Frankish, 1976). Individual lists were compiled by sampling the digits drawn randomly without replacement. Presentation rate was 1000ms; each spoken item had a duration of 750ms and was followed by an ISI of 250ms. The duration of the interpolated processing task was 6000ms, starting after the presentation of the first list item. The processing activity required participants to judge whether pairs of spoken letters rhymed. The letter pairs consisted of monosyllabic English alphabet letter names. Pairs were constrained to avoid successive letters in the alphabet (e.g. J,K), highly confusable fricative letter names (e.g. F,S), and familiar acronyms (e.g. PC, IT, GB). Each letter sound was presented within a 800ms window, followed by a 200ms window of silence before the onset of the second letter sound. The task was participant-paced with an inter-stimulus interval (ISI) of 200ms between a response and the onset of the next letter pair. New letter pairs were not presented if there was <500ms remaining of the 6000ms window. Participants were able to respond at the onset of the second letter in a pair by clicking on an on-screen “rhyme” or “non-rhyme” button.

For the visuo-spatial complex span task, participants were required to remember a series of static, visually presented abstract line figures for serial recall. Each stimulus was presented on screen for 1000ms (750ms followed by a 250ms ISI). The stimuli set consisted of 9 line figures presented in random order. A dynamic visual processing task was interpolated between each line figure. This required participants to mentally rotate two identically-shaped polygons to an upright position to decide whether they were pointing in the same or opposite (mirror image) direction. The polygon on the left was always presented in an upright position. The one on the right could appear at one of 7 rotated positions (45 ^°^, 90°, 135 ^°^, 180^°^, 225 ^°^, 270 ^°^, 315^°^). Pairs of polygons were presented in a random order from a choice of 5 shapes (total stimuli set size of 70). The shapes remained on screen until a response was made. An ISI of 200ms followed a response before the onset of the next polygon pair. New pairs were not presented if there was <500ms remaining of the 6000ms window. Participants could respond by clicking an on screen “same” or “mirror” button as soon as each pair of shapes was presented.

##### N-back

Two variants of the n-back task were used – one verbal and one visuo-spatial. For both tasks, participants were presented with a series of stimuli one at a time. The stimuli presented were identical to the storage items used in the complex span transfer tasks (auditory p-centered digits for the verbal *n*-back task, and static visually presented abstract line figures for the visuo-spatial task). Participants were to judge whether each stimuli matched the one presented *n* items previously in the sequence by pressing the down arrow on the keyboard if it was a match (target) and by not responding if it was not a match (non-target). The inter-stimulus interval was 2500ms. Participants could respond as soon as the stimulus had been presented. Stimuli were presented in blocks of 20+*n* trials, with 6 targets and 14+*n* non-targets presented randomly in each block. Errors included misses (failing to respond to a target) and false alarms (responding to a non-target). Both tasks started with two blocks of practice trials at 1-back. Test trials began with a block at 1-back. If there were fewer than five errors (sum of misses and false alarms) the level of *n* increased by one in the next block. If there were five or more errors the tasks discontinued and the maximum *n*-level reached to this point was scored.

#### Training tasks

##### Complex span

Participants in the complex span training group completed 32 trials per training session, 16 trials each on the verbal and visuo-spatial tasks. The training tasks had the same task structure as the transfer tasks with storage items interpolated with a same-domain processing task. The presentation rate of the storage items and overall length of each processing episode (6000ms) was identical to the transfer tasks. To account for learning of the processing items across training sessions, the presentation rate of the stimuli within each processing window was titrated to individual performance for the training tasks. To determine these presentation rates, participants completed 2 minutes of the processing task before each training session. Average RTs for correct items were calculated and 50% was added to calculate the rate of presentation during training.

Training started with a list length of one (one storage item, one processing episode) in the first session. Task difficulty was adapted across the 20 training sessions based on individual performance. There was an increase of 1 storage item and corresponding processing episode following 2 consecutive correct trials (a correct trial being the storage item(s) recalled in serial order and accuracy >75% across the processing episodes). Task difficulty decreased by 1 storage item and processing episode if there were 2 consecutive incorrect trials (storage items recalled incorrectly and processing performance <75%). Otherwise, difficulty remained the same. Each new training session started at one span level lower than the level reached at the end of the previous session. The average span level reached in each training session was recorded.

The storage items for the verbal complex span training task were consonants presented visually on-screen. The interpolated processing task was a lexical decision task. Words and nonwords were presented one at a time visually on-screen. Participants were required to judge whether each stimulus was a real word by clicking on “word” or “nonword” buttons. They could respond as soon as each stimulus was presented. The word stimuli were drawn at random from a pool of 187 items generated by searching the MRC Psycholinguistic database (Colheart, 1981) for single syllable words of 4-6 letters with Kucera-Francis written word frequency > 50 per million (Kučera & Francis, 1967). An equivalent numbers of nonwords were constructed by blending words from the real word stimuli set (e.g. swapping the initial consonant or consonant cluster) to form nonwords with a similar phonotactic composition.

For the visuo-spatial complex span training task, participants were required to recall a series of spatial locations presented dynamically in a 4×4 grid (akin to a Corsi task). The presentation of each storage item was interleaved with a symmetry decision task that involved judging whether patterns presented on screen were symmetrical. Participants could respond as soon as each pattern was presented by clicking on-screen buttons labeled “symmetrical” and “asymmetrical”.

##### N-back

Participants completed 10 blocks of verbal *n*-back and 10 blocks of visuo-spatial *n*-back in each training session, totalling 20 blocks per session. The n-back training tasks differed from the transfer tasks only in the stimuli – all other features including the presentation rate of the stimuli were the same as in the training tasks. For the verbal *n*-back training, consonants were presented visually on-screen. For the visuo-spatial *n*-back training, spatial locations were presented sequentially one at a time as highlighted green squares in a 4×4 grid. Training started with a block of trials at 1-back and adapted up and down to match participants’ current performance throughout each training session. If there were fewer than three errors (sum of misses and false alarms) in block, the level of *n*-back increased by one in the next block. If there were five or more errors in a block the level of n decreased by one in the subsequent block. In all other cases the level of *n* remained the same. Each new training session started at *n*-1 from the end of the previous training session. The average level of *n* reached in each session was scored.

### Analysis Plan

Bayesian ANOVAs were conducted to analyse on-task training gains across the four training tasks with session (2 to 20) and training task (four training tasks) entered as factors. To investigate whether transfer effects differed by group, Bayesian general linear regression models (GLMs) were performed separately for each transfer task. As there were three groups, three dummy variables were used to run two regression models for each outcome variable. In the first, dummy variables one (D1) and two (D2) were entered to compare the effects of *n*-back training to complex span training (D1 result) and to compare complex span training to no intervention (D2 result). To compare the effects of *n*-back training to the no intervention group, a second regression model was run for each measure in which D2 and dummy variable three (D3) were entered (D2 result). The outcomes of these models also replicated the comparison between the *n*-back and complex span training groups (D3 result) obtained in the first regression models.

Three questions were addressed in the analyses. The first is whether the outcome measures improve after training for each group. The second is whether there is evidence for transfer to each outcome measure for the n-back and complex span training groups relative to the no intervention group. The final question is whether there is evidence for each training condition relative to the other paradigm (complex span or n-back). This is the most stringent comparison that allows us to examine the paradigm-specificity of transfer.

All results are reported as BF. Bayesian methods were conducted in JASP (Love, Selker, Marsman, Jamil, Verhagen & Ly, 2015) with default prior scales. Inverse Bayes Factors (BF_10_) are used to express the odds in favour of the alternative hypothesis compared to the null (Jeffreys, 1961). Values lower than .33 provide evidence for the null hypothesis (no training or transfer), values .33-3 provide equivocal evidence for both hypotheses, and those higher than 3 provide evidence in favour of the alternative hypothesis (training and transfer effects).

### Procedure

Participants were pseudo-randomly assigned to complex span training, n-back training or no training at recruitment (*n*=16 per group). Stratified randomization was used to ensure the groups were matched at baseline in terms of age and gender. All participants completed four tasks during a Time 1 assessment at the MRC Cognition & Brain Sciences Unit lasting approximately one hour. Participants assigned to either of the active training groups also completed their first training session during this visit, adding another 30 minutes to the session. Trainees completed the subsequent 18 online training sessions remotely (e.g. at home) before returning to the MRC Cognition & Brain Sciences Unit to complete their final training session and a Time 2 assessment that was identical to the Time 1 assessment. Participants in the no training group were contacted and asked to return to the MRC Cognition and Brain Sciences Unit for their Time 2 assessment at intervals equivalent to those taken for the trainees. There was substantial evidence for a null effect for differences in the intervals between Time 1 and Time 2 between groups, BF_10_=.185. Participants were paid £6 per hour for their time plus a contribution towards travel costs. All tasks were hosted by the online platform Cambridge Brain Sciences (www.cambridgebrainsciences.com).

Training task order was counterbalanced across participants with half of the participants completing the verbal then the visuo-spatial task in each session, and half completing the visuo-spatial then the verbal task in each session. All participants completed 20 training sessions, but the data failed to upload for the final two sessions for five participants in the complex span training group. Analyses testing for order effects were run including only sessions with complete data for all participants (Sessions 1 to 18 for both tasks). There was equivocal evidence for order effects for gains on the verbal complex span, BF_10_ =2.458 and the visuo-spatial complex span training tasks, BF_10_ =1.478.

Data failed to upload for the final two sessions for three participants on the n-back training tasks. Analyses testing for the effect of training task order were run including sessions where there was complete data for all participants (Sessions 2 to 18 for both tasks). There was equivocal evidence for an order effect for the verbal *n*-back, BF_10_ =2.435 and the visuo-spatial n-back training tasks, BF_10_ =.48.

## Results

### Training task progress

To provide a comparable measure of changes across time across each training task, a standard gain score was calculated by dividing the difference between the average difficulty level attained on each session and the first session by the SD for the first session (Harrison et al., 2013). These scores relative to the first session of training are shown in Figure 2. Improvements of at least 1SD from baseline were observed on all training tasks.

**Figure 2.**
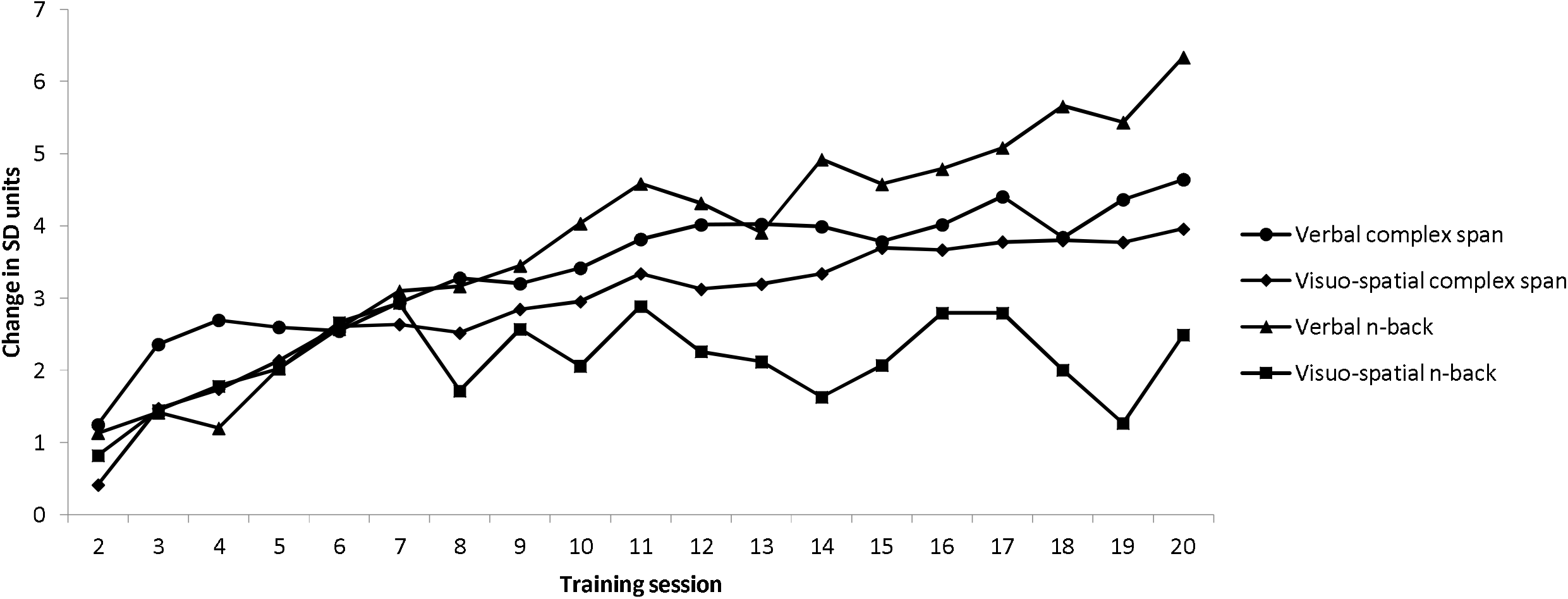
Average difficulty level attained on each session was converted to standard deviation units relative to session 1 (e.g. (session 2 - session 1) / SD session 1; (session 3 – session 1)/session 1 SD. Note that averages are based on the maximum number of data points available for that session.

Descriptive statistics for the four training tasks are reported in Table 2. Note that Session 1 was not included as participants were reaching baseline during this session. Gains on the four training tasks were examined in Bayesian ANOVAs performed on the average difficulty levels of each session from 2 to 20. A 4 x 19 Bayesian repeated measures ANOVA with session (2 to 20) and training task (four training tasks) provided evidence for main effects of session, BF_10_= >100 and condition BF_10_= >100. There was substantial evidence for an interaction between task and session, BF_10_=>100. Post hoc analyses revealed that gains were greater on the verbal complex span training task relative to both the visuo-spatial complex span and the visuo-spatial n-back training tasks.

**Table 2.**
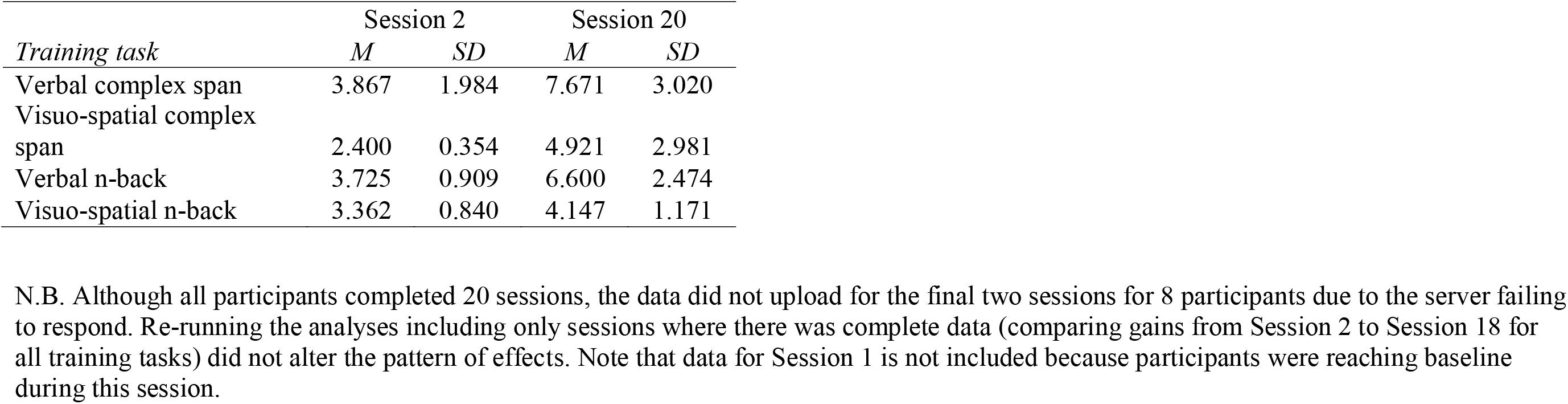
On-task training gains

### Transfer

Bayesian t-tests compared pre- to post-scores for each of the four outcome measures for each training group (see Table 3). There was equivocal evidence for transfer to verbal complex span, verbal n-back and visuo-spatial n-back following complex span training, with evidence favouring a null effect for transfer to visuo-spatial complex span following complex span training (BF=.325). There was strong evidence for transfer for the *n*-back training group on both the verbal and visuo-spatial *n*-back transfer tasks (BF= 23.18 for verbal and BF=20.02 for visuo-spatial n-back transfer tasks). Evidence favoured a null effect for transfer to the complex span outcome measures following n-back training (BF=.33 and .34 for verbal and visuo-spatial complex span respectively). There was equivocal evidence for both the null and alternative hypotheses across the four transfer tasks for the no intervention group (BFs .35 as above to 1).

**Table 3.**
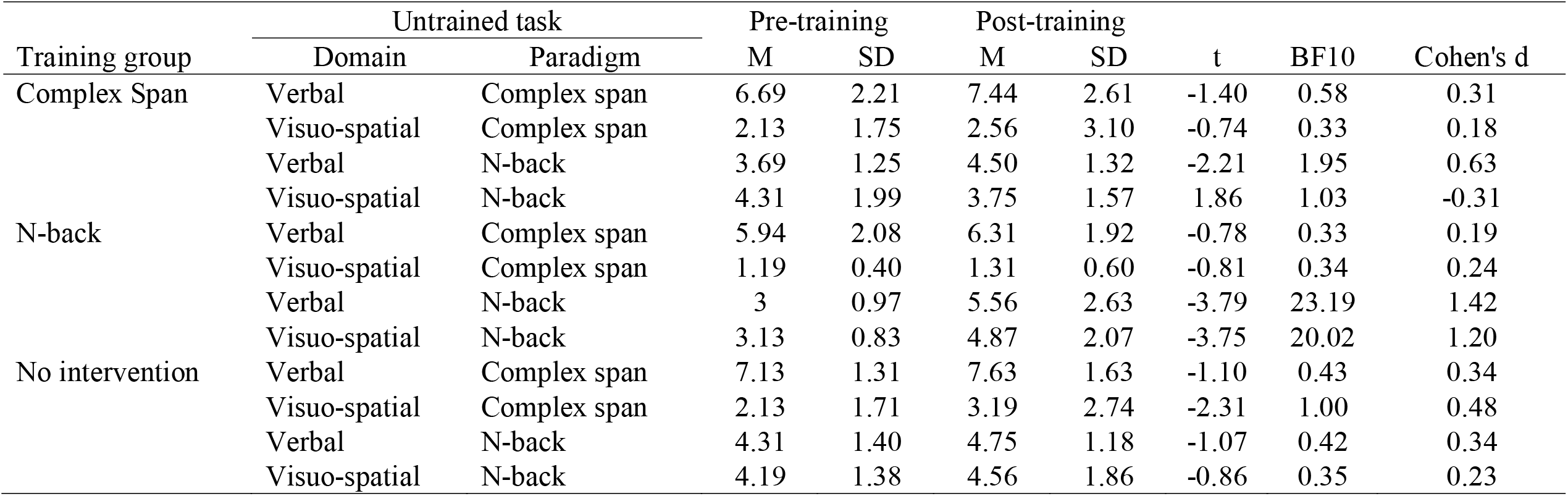
Within group transfer to untrained tasks

| Training group | Untrained task |  | Pre-training |  | Post-training |  | t | BF10 | Cohen's d |
| --- | --- | --- | --- | --- | --- | --- | --- | --- | --- |
|  | Domain | Paradigm | M | SD | M | SD |  |  |  |
| Complex Span | Verbal | Complex span | 6.69 | 2.21 | 7.44 | 2.61 | -1.40 | 0.58 | 0.31 |
|  | Visuo-spatial | Complex span | 2.13 | 1.75 | 2.56 | 3.10 | -0.74 | 0.33 | 0.18 |
|  | Verbal | N-back | 3.69 | 1.25 | 4.50 | 1.32 | -2.21 | 1.95 | 0.63 |
|  | Visuo-spatial | N-back | 4.31 | 1.99 | 3.75 | 1.57 | 1.86 | 1.03 | -0.31 |
| N-back | Verbal | Complex span | 5.94 | 2.08 | 6.31 | 1.92 | -0.78 | 0.33 | 0.19 |
|  | Visuo-spatial | Complex span | 1.19 | 0.40 | 1.31 | 0.60 | -0.81 | 0.34 | 0.24 |
|  | Verbal | N-back | 3 | 0.97 | 5.56 | 2.63 | -3.79 | 23.19 | 1.42 |
|  | Visuo-spatial | N-back | 3.13 | 0.83 | 4.87 | 2.07 | -3.75 | 20.02 | 1.20 |
| No intervention | Verbal | Complex span | 7.13 | 1.31 | 7.63 | 1.63 | -1.10 | 0.43 | 0.34 |
|  | Visuo-spatial | Complex span | 2.13 | 1.71 | 3.19 | 2.74 | -2.31 | 1.00 | 0.48 |
|  | Verbal | N-back | 4.31 | 1.40 | 4.75 | 1.18 | -1.07 | 0.42 | 0.34 |
|  | Visuo-spatial | N-back | 4.19 | 1.38 | 4.56 | 1.86 | -0.86 | 0.35 | 0.23 |

The second set of analyses examined evidence for transfer following training relative to the no intervention condition. There was equivocal evidence for group differences at baseline on the verbal complex span, BF_10_ =.478, Cohen’s d=.530, visuo-spatial complex span, BF_10_ =.782, Cohen’s d=.639, and visuo-spatial *n*-back tasks, BF_10_ =1.176. Post-training scores were therefore entered as the dependent variable with group and pre-training scores entered as independent variables in each of the Bayesian linear regressions for these three outcome measures. There was evidence for baseline group differences on the verbal *n*-back task, BF_10_ =3.811, Cohen’s d=.908. To investigate whether group differences in post-test scores on this task were associated with differences in baseline scores both centred baseline scores and centred baseline score x group product terms were also entered into the regression models. Product terms were derived by the product of the centred scores (individual score minus the group mean) and the grouping variable. The product terms were not significantly associated with post-training scores, meaning the general linear regression models could be run without the product term or centred scores in the final analyses (i.e. with post-training scores as the dependent variable and group and pre-training scores as independent variables).

The outcomes of the regression analyses for each transfer task comparing each active training group to the no intervention group (testing for re-test effects) are reported in Table 4. There was substantial evidence for an absence of transfer to verbal complex span following complex span training relative to the no intervention group (BF=.291). For the visuo-spatial complex span, verbal and visuo-spatial complex, the outcomes were equivocal (BFs .377 to 1.21). Equivocal outcomes were also observed for the visuo-spatial complex span and verbal and visuo-spatial n-back transfer tasks following n-back training (BFs.392 to 1.032). The evidence favoured the null hypothesis for the verbal complex span transfer task (BF=.327), indicating that the changes on this task following n-back training were no greater than for the no intervention group.

**Table 4.** Transfer for each active training group relative to the no intervention group

| Training group |  | Untrained task |  |  |  |
| --- | --- | --- | --- | --- | --- |
|  |  | Verbal<br>Complex span | Visuo-spatial<br>Complex<br>span | Verbal<br>n-back | Visuo-spatial<br>n-back |
| Complex Span | Beta | -0.01 | 0.12 | 0.02 | 0.23 |
|  | t | -0.10 | 1.00 | 0.13 | 1.64 |
|  | BF<sub>10</sub> | 0.29 | 0.38 | 1.21 | 0.89 |
|  | Cohen's d | 0.03 | 0.25 | 0.05 | 0.48 |
| N-back | Beta | 0.14 | 0.15 | -0.30 | -0.26 |
|  | t | 0.96 | 1.21 | -1.64 | -1.77 |
|  | BF<sub>10</sub> | 0.33 | 0.39 | 0.55 | 1.03 |
|  | Cohen's d | 0.29 | 0.31 | 0.64 | 0.56 |

The third set of analyses compared transfer to each outcome measures across the two active training groups (see Table 5). There was substantial evidence for the null hypothesis for the visuo-spatial complex span task showing that changes on this transfer measure were not selective to the complex span training group (BF=.287). There was equivocal evidence for the null and alternative hypotheses for the verbal complex span transfer task when the two active training groups were compared. Combined with the outcome of the analyses comparing the complex span training group to the no intervention group on the verbal complex span transfer task (BF=.291), this provides evidence that there is no selective transfer for verbal complex span following complex span training. Evidence for differences in transfer on the verbal n-back transfer task between the n-back and complex span training groups was equivocal (BF=.542), as for equivocal differences between the n-back and no intervention groups on this transfer measure. It is therefore not possible to determine whether there is a selective enhancement to verbal n-back following n-back training. There was substantial evidence for a group effect for the visuo-spatial *n*-back transfer task, with greater gains found for those who trained on *n*-back relative to those who trained on complex span (BF_10_=9.702).

**Table 5.**
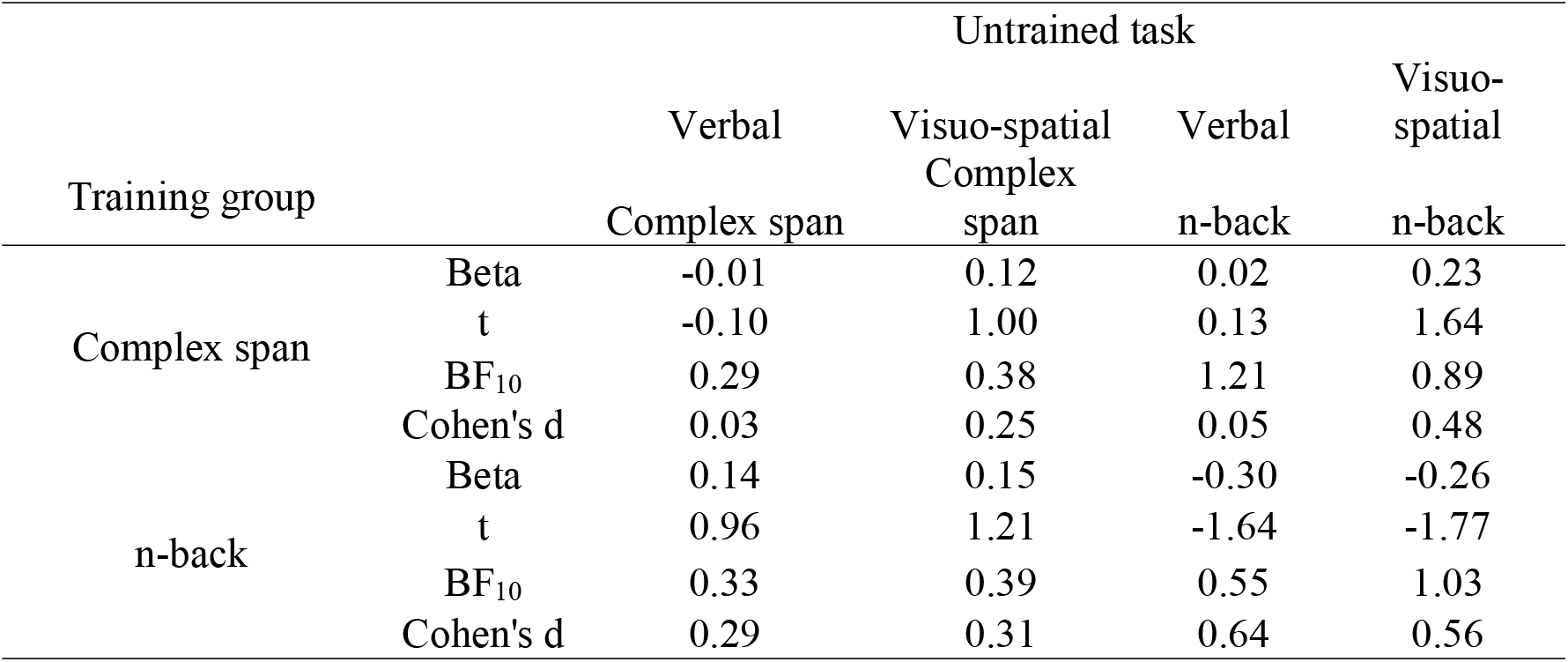
Selective transfer relative to the other active training condition

| Training group |  | Untrained task |  |  |  |
| --- | --- | --- | --- | --- | --- |
|  |  | Verbal<br>Complex span | Visuo-spatial<br>Complex<br>span | Verbal<br>n-back | Visuo-spatial<br>n-back |
| Complex span | Beta | -0.01 | 0.12 | 0.02 | 0.23 |
|  | t | -0.10 | 1.00 | 0.13 | 1.64 |
|  | BF<sub>10</sub> | 0.29 | 0.38 | 1.21 | 0.89 |
|  | Cohen's d | 0.03 | 0.25 | 0.05 | 0.48 |
| n-back | Beta | 0.14 | 0.15 | -0.30 | -0.26 |
|  | t | 0.96 | 1.21 | -1.64 | -1.77 |
|  | BF<sub>10</sub> | 0.33 | 0.39 | 0.55 | 1.03 |
|  | Cohen's d | 0.29 | 0.31 | 0.64 | 0.56 |

## Discussion

This experiment compared transfer patterns from two working memory training paradigms (n-back and complex span) matched for memory items (letters and spatial locations) and training intensity to outcome measures that were also matched for memory items (digits and objects). There was no evidence for transfer across the two paradigms under these matched conditions. This is consistent with many previous studies in which less systematic designs have been used to map transfer (Bürki et al., 2014; Chooi & Thompson, 2012; Jaeggi et al., 2008; Jaeggi et al., 2010; Li et al., 2008; Lilienthal et al., 2013; Minear et al., 2016; Oelhafem, Nikolaidis, Padovani, Blaser, Koenig, & Perrig, 2013; Redick et al., 2013; Schwarb et al., 2015; Sprenger et al., 2013; von Bastian et al., 2013).

This lack of cross-paradigm transfer is consistent with theories that explain training gains in terms of working memory processes specific to particular paradigms.Dahlin et al. (2008), for example, proposed that updating the contents of working memory is trainable. This process would not be expected to contribute to non-updating paradigms such as complex span. The findings also fit with the suggestion that training involves learning (in the form of the development of novel cognitive routines) how to perform specific working memory tasks (Gathercole et al., 2018). The coordinated set of processes necessary to accomplish n-back (in which the greatest challenge is to keep track of the last n items) and complex span (requiring maintenance of stimulus items in the face of distraction) have little in common and hence, as indeed found here, to show little transfer. Note, though, that the present findings cannot distinguish between process- and paradigm specific transfer boundaries. To do this it would be necessary to track transfer across two paradigms sharing a common working memory process, such as running span and n-back updating tasks. Note that although Dahlin et al. (2008) did this and found improvements in the untrained task, the study lacked an active control condition.

In line with previous findings, there was evidence suggesting n-back training transfers to n-back tasks with new stimulus materials (Anguera et al., 2012; Jaeggi et al., 2010; Küper & Karbach, 2015; Minear et al., 2016; Soveri et al., 2017). There was strong evidence for substantial improvements in the two untrained n-back tasks containing digits and visual objects following training on n-back activities with spatial locations and letters. Relative to a no intervention group, there was no strong evidence for the benefits of n-back training for untrained n-back tasks. Compared with complex span training there was substantial evidence that n-back training led to greater improvements on the visuo-spatial n-back task. This suggests indicates that material specificity is not a boundary condition to transfer with n-back training. Instead, training-related changes (possibly in the formation of a new cognitive routine for n-back) appears to operate at a more general level than item-level category.

The strength of transfer from complex span training to other complex span tasks with different materials and distractor processing was more equivocal, varying between weak or absent. Performance on the untrained visuo-spatial complex span task did not improve following complex span compared with n-back training. There was also no evidence for greater gains for the complex span training group on the verbal complex span transfer task relative to the test re-test group. This suggests that the cognitive processes modified during training on complex span are tied to the training task materials (either the memory items or the distractor stimuli). This outcome has implications for the nature of the putative cognitive routines developed during training (Gathercole et al., 2018). A critical feature of a successful routine for complex span could be coordinated processes designed to maintain stimulus items while performing interpolated distraction activities. This might, for example, involve rapid switching between the distractor activity and either rehearsal of the memory items (Towse, Hitch & Hutton, 1999) or refreshing them attentionally (Barrouillet, Bernadin & Camos, 2004). The present findings suggest that a sub-routine that controls such a process is likely to be specific the particular memory items and distractor activities engaged by the training task: it does not operate at a more general level that would generalize across these changes in materials. There is little evidence that participants receiving complex span training in this experiment do not appear to have learned at a general level how to carry out the attentional shifting or other types of processes to protect memory representations from interference. Instead, they appeared to learn how to do so in the context of the specific trained tasks.

The present findings both add to other evidence for lack of transfer across complex span tasks (Blacker et al., 2017; Minear et al., 2016) and stand in contrast to other reports of transfer across different memory items and interpolated distraction activities (e.g. Harrison et al., 2013; Henry et al., 2014; Richmond et al., 2014; von Bastian et al., 2013). Why the outcomes are so variable across studies requires further systematic investigation.

In summary, the current findings provide evidence that the cognitive changes resulting from intensive working memory training are relatively specific not only to the training paradigms but, in the case of complex span, to the detailed characteristics of the distractor activities. The data clearly establish that transfer does not extend across global changes in working memory paradigm and is even limited within task. While the benefits of training extend to n-back tasks employing other stimuli, for complex working memory tasks they are much more restricted. Any potential learning of the unfamiliar task during training thus appears to be highly specific.

## Author note

JH and SG led the conception and design of the work and took primary responsibility for drafting the manuscript. FW collected the data, and AH developed the online tasks. FW and AH commented on drafts. Correspondence concerning this article should be sent to Joni Holmes. This research was supported by the Medical Research Council of Great Britain, the University of Cambridge.

## References

Anguera, J. A., Bernard, J. A., Jaeggi, S. M., Buschkuehl, M., Benson, B. L., Jennett, S., … & Seidler, R. D. (2012). The effects of working memory resource depletion and training on sensorimotor adaptation. Behavioural Brain Research, 228(1), 107-115.

Barrouillet, P., Bernardin, S., & Camos, V. (2004). Time constraints and resource sharing in adults’ working memory spans. Journal of Experimental Psychology: General, 133(1), 83.

Blacker, K. J., Negoita, S., Ewen, J. B., & Courtney, S. M. (2017). N-back versus complex span working memory training. Journal of cognitive enhancement, 1(4), 434-454.

Bürki, C. N., Ludwig, C., Chicherio, C., & de Ribaupierre, A. (2014). Individual differences in cognitive plasticity: an investigation of training curves in younger and older adults. Psychological Research, 78(6), 821-835.

Chooi, W. T., & Thompson, L. A. (2012). Working memory training does not improve intelligence in healthy young adults. Intelligence, 40(6), 531-542.

Coltheart, M. (1981). The MRC psycholinguistic database. The Quarterly Journal of Experimental Psychology, 33(4), 497-505.

Dahlin, E., Neely, A. S., Larsson, A., Backman, L., & Nyberg, L. (2008). Transfer of learning after updating training mediated by the striatum. Science, 320(5882), 1510-1512.

De Simoni, C., & von Bastian, C. C. (2018). Working memory updating and binding training: Bayesian evidence supporting the absence of transfer. Journal of Experimental Psychology: General, 147(6), 829.

Dunning, D. L., & Holmes, J. (2014). Does working memory training promote the use of strategies on untrained working memory tasks? Memory and Cognition, 42(6), 854-862.

Gathercole, S.E., Dunning, D.L., Holmes, J. & Norris, D. (2018). Working memory training involves learning new skills, under revision.

Harrison, T. L., Shipstead, Z., Hicks, K. L., Hambrick, D. Z., Redick, T. S., & Engle, R. W. (2013). Working memory training may increase working memory capacity but not fluid intelligence. Psychological Science, 24(12), 2409-2419.

Henry, L. A., Messer, D. J., & Nash, G. (2014). Testing for near and far transfer effects with a short, face-to-face adaptive working memory training intervention in typical children. Infant and Child Development, 23(1), 84-103.

Jaeggi, S. M., Buschkuehl, M., Jonides, J., & Perrig, W. J. (2008). Improving fluid intelligence with training on working memory. Proceedings of the National Academy of Sciences of the United States of America, 105(19), 6829-6833.

Jaeggi, S. M., Buschkuehl, M., Perrig, W. J., & Meier, B. (2010). The concurrent validity of the N-back task as a working memory measure. Memory, 18(4), 394-412.

Jeffreys, H. (1961). Theory of probability. Oxford, UK: Open University Press.

Kane, M. J., Conway, A. R., Miura, T. K., & Colflesh, G. J. (2007). Working memory, attention control, and the N-back task: a question of construct validity. Journal of Experimental Psychology: Learning, Memory, & Cognition, 33(3), 615.

Kučera, H., & Francis, W. N. (1967). Computational analysis of present-day American English. Dartmouth Publishing Group.

Küper, K., & Karbach, J. (2016). Increased training complexity reduces the effectiveness of brief working memory training: evidence from short-term single and dual n-back training interventions. Journal of Cognitive Psychology, 28(2), 199-208.

Li, S. C., Schmiedek, F., Huxhold, O., Röcke, C., Smith, J., & Lindenberger, U. (2008). Working memory plasticity in old age: practice gain, transfer, and maintenance. Psychology & Aging, 23(4), 731.

Lilienthal, L., Tamez, E., Shelton, J. T., Myerson, J., & Hale, S. (2013). Dual n-back training increases the capacity of the focus of attention. Psychonomic Bulletin & Review, 20(1), 135-141.

Love, J., Selker, R., Marsman, M., Jamil, T., Verhagen, A. J., & Ly, A. (2015). JASP (Version 0.6. 6). Computer software.

Minear, M., Brasher, F., Guerrero, C. B., Brasher, M., Moore, A., & Sukeena, J. (2016). A simultaneous examination of two forms of working memory training: Evidence for near transfer only. Memory & Cognition, 44(7), 1014-1037.

Morton, J., Marcus, S., & Frankish, C. (1976). Perceptual centers (P-centers). Psychological Review, 83(5), 405.

Oelhafen, S., Nikolaidis, A., Padovani, T., Blaser, D., Koenig, T., & Perrig, W. J. (2013). Increased parietal activity after training of interference control. Neuropsychologia, 51(13), 2781-2790.

Redick, T. S., Shipstead, Z., Harrison, T. L., Hicks, K. L., Fried, D. E., Hambrick, D. Z., … & Engle, R. W. (2013). No evidence of intelligence improvement after working memory training: a randomized, placebo-controlled study. Journal of Experimental Psychology: General, 142(2), 359.

Richmond, L. L., Wolk, D., Chein, J., & Olson, I. R. (2014). Transcranial Direct Current Stimulation Enhances Verbal Working Memory Training Performance over Time and Near Transfer Outcomes. Journal of Cognitive Neuroscience, 26(11), 2443-2454.

Schwarb, H., Nail, J., & Schumacher, E. H. (2016). Working memory training improves visual short-term memory capacity. Psychological Research, 80(1), 128-148.

Soveri, A., Antfolk, J., Karlsson, L., Salo, B., & Laine, M. (2017). Working memory training revisited: A multi-level meta-analysis of n-back training studies. Psychonomic Bulletin & Review, 1-20.

Sprenger, A. M., Atkins, S. M., Bolger, D. J., Harbison, J. I., Novick, J. M., Chrabaszcz, J. S.,… Dougherty, M. R. (2013). Training working memory: Limits of transfer. Intelligence, 41(5), 638-663.

Thompson, T. W., Waskom, M. L., Garel, K. L. A., Cardenas-Iniguez, C., Reynolds, G. O., Winter, R., … & Gabrieli, J. D. (2013). Failure of working memory training to enhance cognition or intelligence. PloS One, 8(5), e63614.

Towse, J. N., Hitch, G. J., & Hutton, U. (1999). The resource king is dead! Long live the resource king! Behavioral & Brain Sciences, 22(1), 111-111.

von Bastian, C. C., Langer, N., Jancke, L., & Oberauer, K. (2013). Effects of working memory training in young and old adults. Memory & Cognition, 41(4), 611-624.

von Bastian, C. C., & Oberauer, K. (2013). Distinct transfer effects of training different facets of working memory capacity. Journal of Memory and Language, 69(1), 36-58.

von Bastian, C. C., & Oberauer, K. (2014). Effects and mechanisms of working memory training: a review. Psychological Research, 78(6), 803-820.

Waris, O., Soveri, A., & Laine, M. (2015). Transfer after working memory updating training. PloS One, 10(9), e0138734.

